# Single-cell quantitation of histones and histone post-translational modifications across the cell cycle by high-throughput imaging

**DOI:** 10.1101/070219

**Authors:** Linda Zane, Fleur Chapus, Gianluca Pegoraro, Tom Misteli

## Abstract

Post-translational modifications (PTM) of histone proteins are critical determinants of genome function. Analysis of histone PTM levels has traditionally been performed using biochemical bulk methods to measure individual modifications. These approaches are generally unable to detect differences in histone and histone PTM levels during the cell cycle, do not provide information on variability amongst individual cells, and they are not suitable for multiplexing to comprehensively analyze large numbers of histone modifications in single experiments. We have developed HiHiMap (**H**igh-throughput **H**istone **Map**ping), an automated high-throughput immunofluorescence technique to determine histone and histone PTM levels across the cell cycle at the single-cell level in a highly parallel format. Our approach uses imaging-based quantification of DNA content and cyclin A levels to stage individual cells in the cell cycle combined with determination of the corresponding histone and histone PMT levels in the same cells. We have applied HiHiMap to a set of 22 histone variants and histone modifications. As proof-of-principle for a biological application, we use HiHiMap to the histone and histone PTMs landscape in primary, immortalized and oncogenically transformed cells. We find differences in the behavior of a specific set of histone PTM during the the cell cycle in transformed cells compared to normal and immortalized cells. The method developed here is widely applicable to the systematic study of histone modifications in physiological and pathological settings at the single-cell level.

## INTRODUCTION

Histones are among the most evolutionarily conserved proteins in eukaryotic cells. Two copies of each core histone (H2A, H2B, H3 and H4) are assembled into octamers around which 147 base pairs of DNA are wrapped to form nucleosomes, the basic structural units of chromatin^1^. Histones are extensively modified on their N-terminal tails by post-translational modifications (PTMs), including phosphorylation on serine or threonine residues, methylation on lysine or arginine, acetylation, ubiquitylation and SUMOylation^2,3^. These modifications exert biological functions by affecting the recruitment of histone binding proteins, such as chromatin remodelers or chromatin structural proteins^4–6^. While some histone PTMs are known to be associated with active chromatin states, others are indicative of repressive chromatin^7^,^8^. In addition to histone PTMs, histone variants also play an important role in the regulation of chromatin organization, for example, the replacement of canonical histones by non-allelic variants modifies nucleosome stability and chromatin dynamics^9^. Histone variants and PTMs are important for regulation of basic genomic processes including the DNA damage response (DDR), transcription and replication^10–14^. Due to their role in gene regulation and genome stability, histone variants and modifications have been implicated in carcinogenesis^15–19^.

To maintain sufficient levels of histones to ensure normal chromosomal DNA compaction throughout the cell cycle, the synthesis of DNA and of core histones is tightly coordinated. Core histone protein levels double via transcriptional upregulation during the cell cycle as the genome is duplicated in S-phase^20–23^. PTMs are generally absent on newly synthesized histones but are re-established upon DNA replication^24–27^. In contrast to the histone proteins themselves, histone PTMs turn over rapidly via action of histone modifying enzymes and may thus fluctuate during the cell cycle, within a cell cycle phase and between individual cells.

The standard methods to analyze histones and histone PTMs are western blotting or stable isotope labeling with amino acids in cell culture (SILAC)-based quantitative mass spectrometry^28 25,29–32^. These methods are limited in that they are only able to provide a population-based average measurement of histone levels or PTMs and are unable to detect differences amongst subpopulations or individual cells. While they can be used for analysis of cell-cycle fluctuations when combined with cell cycle synchronization methods, the synchronization and labeling steps require extensive culturing of cells over several days, relatively large numbers of cells, and are not suitable for parallel analysis of large numbers of modifications.

To overcome these limitations, we developed HiHiMap (**Hi**gh-throughput **Hi**stone **Map**ping), a method for single-cell quantification of histone and histone PTM levels throughout the cell cycle. The approach is based on high-throughput imaging and uses cyclin A immunofluorescence (IF) and DAPI staining for cell cycle staging which allows the accurate determination of the cell cycle stage of individual cells and, simultaneously, the measurement of histone and histone PTM levels at the single-cell level. The high-throughput nature of the method allows for parallel analysis of multiple histone PTMs in a single experiment. Using HiHIMap, we examined 22 histone and histone modifications in an oncogenic cellular transformation model^33–36^ and identified single cell-level changes of several histone variants and PTMs during oncogenic transformation.

## MATERIALS AND METHODS

### Cell culture

hTERT-immortalized human diploid fibroblasts (HDFs), CRL-1575, and primary HDFs from healthy Caucasian individuals (AG06310A, AG06289A and AG04551A) (Coriell Cell Repository) were cultured in Minimum Essential Medium (MEM from Life Technologies) supplemented with 15% (v/v) Fetal Bovine Serum (FBS, Atlanta Biologicals), 1% (v/v) Penicillin/Streptomycin (P/S) and 1% (v/v) L-Glutamine (Life Technologies).

### Cell transformation

Immortalized and transformed human cell lines were generated as described^33^ from HDFs. Recombinant retroviral vectors expressing human telomerase hTERT, SV40 early region (LT and ST antigens) and oncogenic H-RasV12 were introduced simultaneously to generate transformed cells. Retroviral vector supernatants were produced by co-transfecting 10-cm dishes of HEK293T cells (ATCC) with each of the packaging plasmid pGAG-pol and pVSV-G and with pBABE-puro-hTERT, pBABE-zeo-large T genomic or pBABE-neo-rasV12 (X-treme GENE HP DNA transfection reagent, Roche). The medium of 293T cells was changed 12 h after transfection and 6 mL per 10-cm dish of fresh medium was added. Viral vector supernatants were harvested 24 h later and used to transduce HDFs with 5 μg/mL polybrene. Typically, 30%-60% transduction efficiency of the HDFs was achieved by using this protocol as measured by parallel transductions with a GFP-expressing retroviral vector instead of the infection with hTERT-expressing retroviral construct. Then, transduced cells were selected using puromycin (5 μg/mL), zeocin (100 μg/mL) or neomycin (0.5 μg/mL) for 15 days to purify polyclonal-transduc ed populations. These immortalized and transformed cell lines were grown as described above. All cells were grown at 37°C in 5% CO_2_.

### Western Blot and qRT-PCR

Briefly, cells were lysed and DNA was sonicated. After determining protein concentration, 5 μg of total proteins was loaded on a NuPAGE™ NovexTM 4-12% Bis-Tris Protein Gel (ThermoFisher). After protein transfer on a PVDF membrane (Immobilon P, Millipore cat#161-0737), expression of SV40LT and H-Ras proteins was measured by incubation of this membrane with SV40LT antibody Ab101 (BD Pharmingen #554149, 1/1000), Pan-Ras antibody (Calbiochem #OP40 - final concentration 0.3 μg/mL). Intensities of protein bands, revealed by chemiluminescent reagents, were measured using a ChemiDocTM MP imaging system from BioRad. The expression of Pan-Ras and SV40LT was normalized to the endogenous expression of GAPDH.

For real-time quantitative reverse transcriptase-polymerase chain reaction (qRT-PCR), total cellular RNA was isolated using the RNeasy kit according to the manufacturer's instructions (Qiagen, cat#74136). The first-strand cDNA was synthesized from 0.5 μg RNA using random primers and MultiScribe reverse transcriptase (from high capacity cDNA reverse transcription kit #4368813, Applied Biosytems). RNA concentration and purity were determined by UV spectrophotometry (Nanodrop). The primer pairs used were as follows: hTERT-F: 5'GAGCTGACGTGGAAGATGAG3'; hTERT-R: 5'CAGGATCTCCTCACGCAGAC3'; V12hRAS_4F: 5'GCAAGTGTGTGCTCTCCTGA3'; V12hRAS_4R: 5'CTGGCGAATTCCTACAGCTT3'; SV40_LT6_F: 5'TTGGAGGCCTAGGCTTTTG3'; SV40_LT6_R: 5'CAGGCACTCCTTTCAAGACC3')^36^. Each primer pair was designed to generate an amplicon across different exons to avoid genomic amplification. GAPDH was used as the reference gene for the normalization of results (GAPDH-R: 5' ACCCACTCCTCCACCTTTGA3'; GAPDH-F: 5'CTGTTGCTGTAGCCAAATTCGT3'). PCR was performed using iQ SYBR Green Supermix on a CFX96 real-time PCR system (Bio-Rad). A large amount of cDNA was prepared from the control cell line, 66^+++^ cells, a transformed human skin fibroblast cell line that forms colonies in soft agar and tumors in immunocompromised mice^36^. The cDNA was diluted 10 times, aliquoted and used as a calibrator. Serial dilutions of 66^+++^ cDNA were used to determine the PCR efficiency of each primer. For relative quantification and normalization, the comparative Ct (or Eff-DDC) method was used^37^.

### Soft Agar Assay

Soft agar assays were performed in three-layer soft agar in 6-well plates. A bottom layer of 0.6% Agar Noble (Lonza) in MEM was first placed onto 6-cm dishes. 5000 immortalized or transformed HDFs per well were seeded in 0.45% top agar in MEM with FBS, P/S and fungizone (250 μg/mL). Fresh top agar was added after 1.5 weeks, and, after 3 weeks, colonies were stained overnight at 37°C with MTT (Sigma) at a concentration of 10 mg/mL (1/100) in PBS. Colonies were counted by using Image J software.

### High-throughput Microscopy

Cells were plated in 384-well Cell Carrier plates (PerkinElmer, Waltham, MA) at a concentration of 60 cells/μL (3000 cells/well) and incubated for 24 h before immunofluorescence (IF) staining. Cells were fixed in 4% paraformaldehyde (PFA) in PBS for 15 min, washed for 5 min three times with PBS and permeabilized in 0.5% Triton X-100/PBS for 15 min at RT. Cells were then washed for 5 min three times with PBS, blocked for 20 min with 3% BSA in PBS and incubated with the indicated primary rabbit or mouse antibodies at the indicated concentrations (Table S1) and anti-cyclin-A2 antibody (1:125, Abcam #ab16726, clone 6E6) for 30 min at RT, washed for 5 min three times with PBS before a 30-min incubation with appropriate anti-rabbit or anti-mouse secondary antibodies labeled with Alexa488 or Alexa-647 (Life Technologies #A11034 (1:200) and #A31571 (1:200)). In most experiments two histones or histone modifications were detected simultaneously per well. After three 5min-washes with PBS, DNA was counterstained with 10 μg/ mL of 4',6'-diamidino-2-phenylindole (DAPI) for 10 min at RT. Cells were left in 100 μL/ well PBS, and either immediately imaged right away or stored at 4°C in sealed plates until imaging.

### Image Acquisition

Imaging was performed on an Opera QEHS high-content screening microscopy system (PerkinElmer) running Opera 2.0.1 software using a 40X water immersion (NA 0.9) or a 20X water immersion (NA 0.7) objective lens and three 12-bit 1.3 Mp CCD cameras with a pixel binning setting of 2, corresponding to a final pixel size of 323 nm for the 40X objective and 646 nm for the 20X.. DAPI, Alexa488 and Alexa647 were sequentially acquired in 150 (when using the 40X lens) or 50 (when using the 20X lens) randomly sampled fields per well using a UV polychrome box (Wide field), a 488 nm laser (Confocal) and a 640 nm laser (Confocal). Typically, at least 3000 cells were imaged for each condition (~1500 cells/well and 2 wells per histone or PTM).

### Automated Image analysis

Acquired images were analyzed with Columbus 2.5 or 2.6 (PerkinElmer) software. Nuclei were detected and non-nuclear objects representing nuclear debris and/or nuclear segmentation errors were eliminated by manually setting nuclear area and roundness filters. The resulting nucleus region of interest was then used as the search region to measure integrated and mean fluorescence intensities of DAPI, cyclin A and histone or PTM of interest in the UV, Alexa 488 and Alexa 647 channels, respectively.

Single cell image analysis results were exported as tab separated text files and further analyzed with R statistical analysis software (https://cran.r-project.org/). Cell cycle phases G1, S and G2/M were determined as described previously^38^ using the DAPI intensity levels. In addition, we identified G2 cells by the marker protein cyclin A2.

### Cell cycle profiling by imaging

For the high-throughput microscopy pipeline, a dedicated R script was generated. First, the cell cycle profiles for each well based on the DAPI integrated intensity were plotted and, after a visualization of the range of G1 and G2 peak values in different wells, the positions of the peak maxima were calculated using a non-parametric mixed model estimation in the R mixtools package^39^ or by manually determining the positions of the peaks when the model was unable to calculate them. The G1 peak value was set to 1 in order to define the same thresholds for G1, S and G2/M phases in all samples (Fig.S1). After plotting a histogram for the cyclin A intensity values, the cyclin A threshold for all wells of a 384-well plate was manually defined based on the background staining in the wells incubated without cyclin A antibody but with the secondary antibodies. Then, the cell cycle stage associated with each individual cell in the dataset was determined based on both cyclin A and DAPI thresholds. Boxplots overlaid with dot plots of the integrated intensity of a histone or PTM at each cell cycle phase were created and text files containing the geometric mean integrated intensity and mean fluorescence intensity for each histone and PTM studied at each cell cycle stage were generated.

To compare the levels of histones and histone PTMs in primary, immortalized and transformed cells, several steps were added to the R script. To take into consideration the ploidy status of the transformed cells, the histone and histone PTM levels were normalized to the DAPI amount. To simplify the representation of the results, we show results for tetraploid cells, which is the major population of transformed cells. For example, the transformed cells of the AG06289 sample displayed a mixed population of diploid and tetraploid cells. The tetraploid cells represent the main population and their histone and histone PTM levels are indicated in the graphs. Note that after normalization to the DNA amount, the results for the tetraploid and diploid cells gave similar results.

### Cell cycle profiling by flow cytometry

Cell cycle distribution was assessed by measuring the DNA content of a suspension of fresh nuclei by flow cytometric analysis after PI staining. Cells from one 60-80% confluent 10 cm-dish were washed with PBS. The supernatant was discarded, and cells were resuspended in 50 μL of PBS and fixed with 450 μL of 70% ethanol for 30 minutes at 4°C and then stored at -20°C. After ethanol elimination, cells were washed twice with PBS and labeled with PI (Sigma #P4864) in the presence of RNase A (Sigma, 100 mg/mL), scanned by flow cytometry using a FACS Calibur and analyzed using BD Cell Quest Pro software.

### Statistical analysis

Bar chart results are represented as means ± 1 SD (standard deviations) of a minimum of three biological replicates. For comparisons of histone and histone PTM levels between groups, results are represented as notched box plots using R^40^ and the ggplot2 package^41^. Box plot notches indicate the estimated 95% confidence interval (CI) for the estimated median value, calculated as 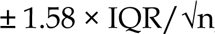, where IQR is the interquartile range or distance between the first and third quartiles, and n is the number of cells^42^. The lower and upper hinges of the boxplots correspond to the first and third quartiles (the 25^th^ and 75^th^ percentiles). The upper and lower whiskers extend from the hinge to ± 1.5 * IQR of the hinge.

The Student’s t test was used for all pairwise comparisons and p-values were adjusted using the Benjamini-Hochberg multiple test correction. P<0.05 was considered statistically significant for these analyses. All statistical analyses were performed using R statistical analysis software.

## RESULTS

### HiHiMap Outline

We sought to develop a method to measure the levels of histone and histone PTMs through the cell cycle at the single-cell level and in a high-throughput format to enable comprehensive analysis of the epigenetic landscape in individual cells. HiHiMap is a high-throughput imaging-based approach in which DAPI and cyclin A IF staining is used to stage individual cells in the cell cycle and this information is combined with the use of antibody-based quantitative detection of histones and histone PTMs in the same individual cells (Fig. 1).

**Figure 1.**
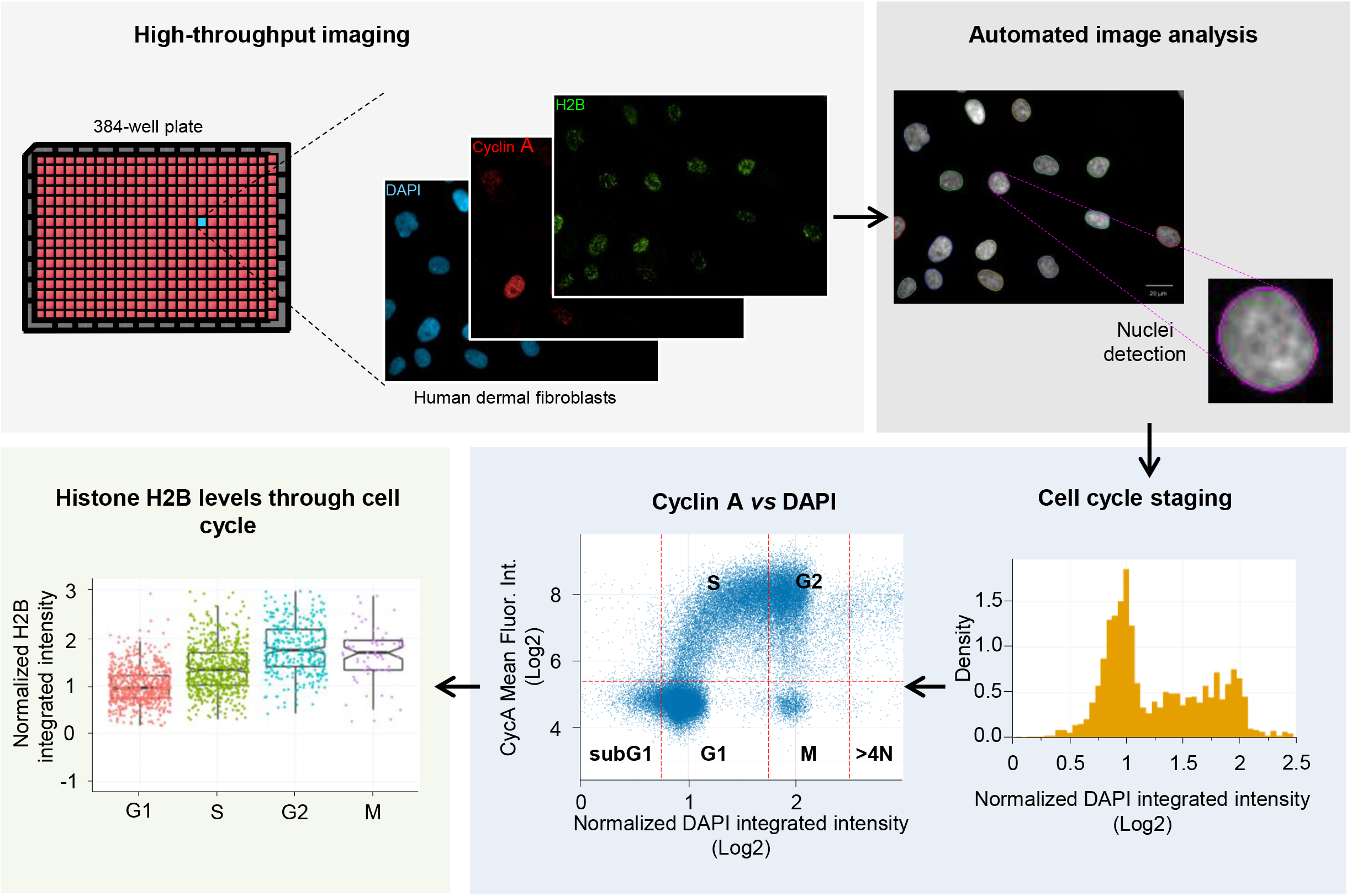
Outline of the high-throughput histone mapping (HiHiMap) method. Cells are plated in 384-well plates, fixed and nuclei are stained with DAPI. Cells are incubated in the presence of an antibody against cyclin A and an antibody against specific histone or histone post-translational modifications. Images of stained cells are acquired in 2D using a high-throughput confocal microscope (Opera, Perkin Elmer) and automated image analysis is used to calculate the averaged integrated and mean fluorescence intensities of DAPI, cyclin A and histone/ PTM of interest in 3 different channels for each cell (Columbus software). The histograms of DNA content in the cell population are then calculated to generate cell cycle profiles using R. The first peak represents the G1 cell population and its position was normalized to a value of 1. Cells in the G2 phase were distinguished from the cells in mitosis using a cyclin A intensity threshold in addition to the DAPI cutoffs. The single-cell levels of normalized fluorescence intensity for a histone or a histone modification is represented for each cell cycle phase. Each dot represents one individual cell.

For HiHiMap, cells are plated in 384-well imaging plates, fixed in 4% paraformaldehyde and stained for cyclin A and a histone or histone PTM of interest for each well using specific antibodies (see Materials and Methods). For analysis, ~ 2, 000 cells in 50 randomly selected fields are imaged per well using a high-throughput microscope (see Materials and Methods). Nuclei are identified by automated image segmentation, and averaged integrated DAPI, mean cyclin A and integrated histone or histone PTM fluorescence intensities are measured in three separate channels (UV, Alexa 488, Alexa 647; see Materials and Methods). Each individual nucleus is assigned to a cell cycle stage based on integrated DAPI intensity and cyclin A staining and the corresponding histone or histone PTM levels are measured in each cell-cycle staged nucleus. Results are represented as single-cell datasets (Fig. 1). As proof-of-principle, HiHiMap was applied to the analysis of 26 histones, histone variants and histone PTMs (Table S1).

### Cell cycle staging

Single cells are staged in the cell cycle by measurement of the integrated DAPI intensity of an individual cell nucleus as previously described^38^. We have previously demonstrated the accuracy of cell cycle staging by DAPI using this approach by direct comparison with FACS analysis^38^. Typical imaging-based DAPI intensity distributions are bi-phasic with peak 1 (arbitrarily defined as intensity value 1; see Methods) representing pre-replicative G1 cells^38^. To unambiguously stage individual cells, DAPI intensity was combined with cyclin A signal, which is known to be elevated in S-and G2-phase^43^. G1 cells were defined as a DAPI intensity of 0.75 - 1.75 and cyclin A negative, S phase cells as DAPI intensity of 0.75 - 1.75 and cyclin A positive, G2 cells as DAPI intensity of 1.75-2.5 and cyclin A positive, and M phase cells as 1.75 - 2.5 and cyclin A negative (Fig. S1).

### HiHiMap validation using core histones

To validate HiHiMap, we measured the level of three nuclear proteins with known cell cycle behavior in hTERT immortalized CRL-1474 human dermal fibroblasts (HDF)^36,44^. Histone H4 is a core histone whose intensity level is known to follow DNA content during the cell cycle^21–23^. As expected, histone H4 levels increase 1.87 ± 0.03 (mean ± SD) and 2.3 ± 0.03-fold in S and G2 phases, respectively, in comparison to G1 phase cells (Fig. 2B and C). Results were highly reproducible in technical and biological replicates (Fig. S2). Phosphorylation on Serine 10 of histone H3 (H3S10Ph) is a well-characterized mitotic marker^45^. As expected, a major increase of H3S10ph levels was found in G2/M phase (9.2 ± 0.7-fold) in comparison to G1 cells (Fig. 2E and F). As a negative control, the transcription factor LHX9, involved in brain development^46^, does not vary with DNA levels in HDF and we only observe a marginal increase in the LHX9 level during the cell cycle (ratios S/G1= 1.2 ± 0.01;G2/G1=1.3 ± 0.02 and M/G1=1.6 ± 0.08) (Fig. 2H and I).

**Figure 2.**
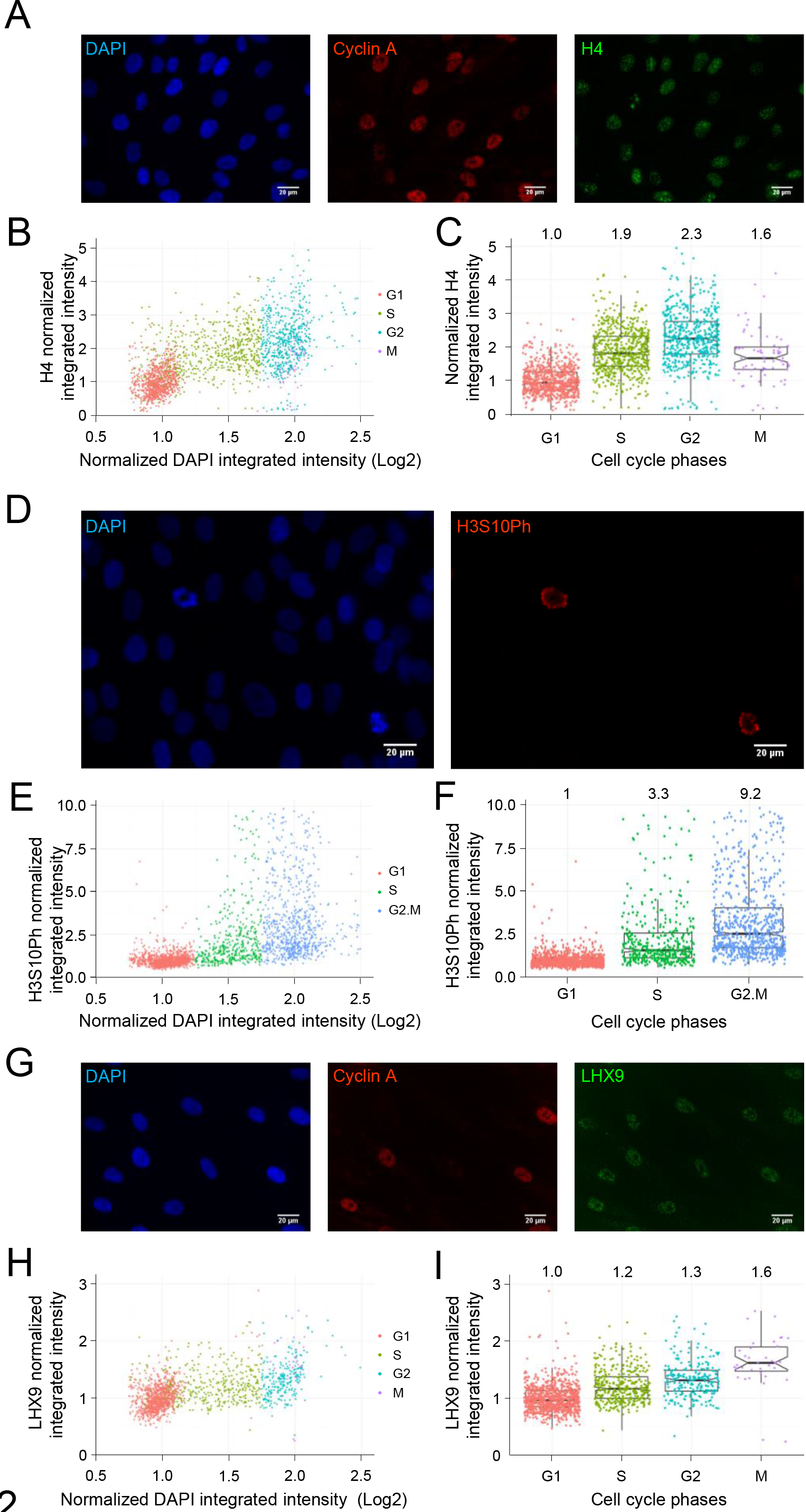
Proof of principle of HiHiMap. Representative confocal images of (A) H4, a core histone, (D) H3S10Ph, a mitotic histone PTM, (G) LHX9, a non-cell cycle regulated transcription factor involved in brain development, and their cyclin A (far red) and/or DAPI staining (blue) in immortalized human dermal fibroblasts. Scale bar, 10 μm. Single-cell levels of H4 (B, C), H3S10Ph (E, F), and LHX9 (H, I) as a function of DNA amount (DAPI intensity level) and at each cell cycle stage. Each dot represents a single cell. In box plots, the lined corresponds to the median, notches represent the estimated 95% confidence interval (CI) for the median, the lower and upper hinges of the box plot indicate the 25th and 75th percentiles and the whiskers correspond to ± 1.5 * IQR of the hinge, where IQR is the interquartile range or distance between the first and third quartiles. The numbers above the box plots represent the means.

### Analysis of core histones and variants during the cell cycle

To extend our analysis of histones, we measured the levels of all core histones (Fig. 3A). Protein levels of H2A, H2B, H3 and H4 increased in S phase by 1.7 ± 0.02-, 1.4 ± 0.02-, 2.0 ± 0.03- and 1.87 ± 0.03-fold, respectively, and 2.3 ± 0.03-, 1.84 ± 0.03-, 2.6 ± 0.04-, 2.3 ± 0.03-fold, respectively, in G2 phase relative to G1 before decreasing 1.7 ± 0.09-, 1.7 ± 0.08-, 1.6 ± 0.09- and 1.7 ± 0.09-fold during mitosis in comparison to G2 phase (Fig. 3A). The 2.6-fold increase in histone H3 level reflects detection of multiple H3 isoforms (H3.1, H3.2 and H3.3) by the antibody used (Fig. 3A). We conclude that HiHiMap accurately detects cell-cycle related fluctuations in histones and histone PTMs.

**Figure 3.**
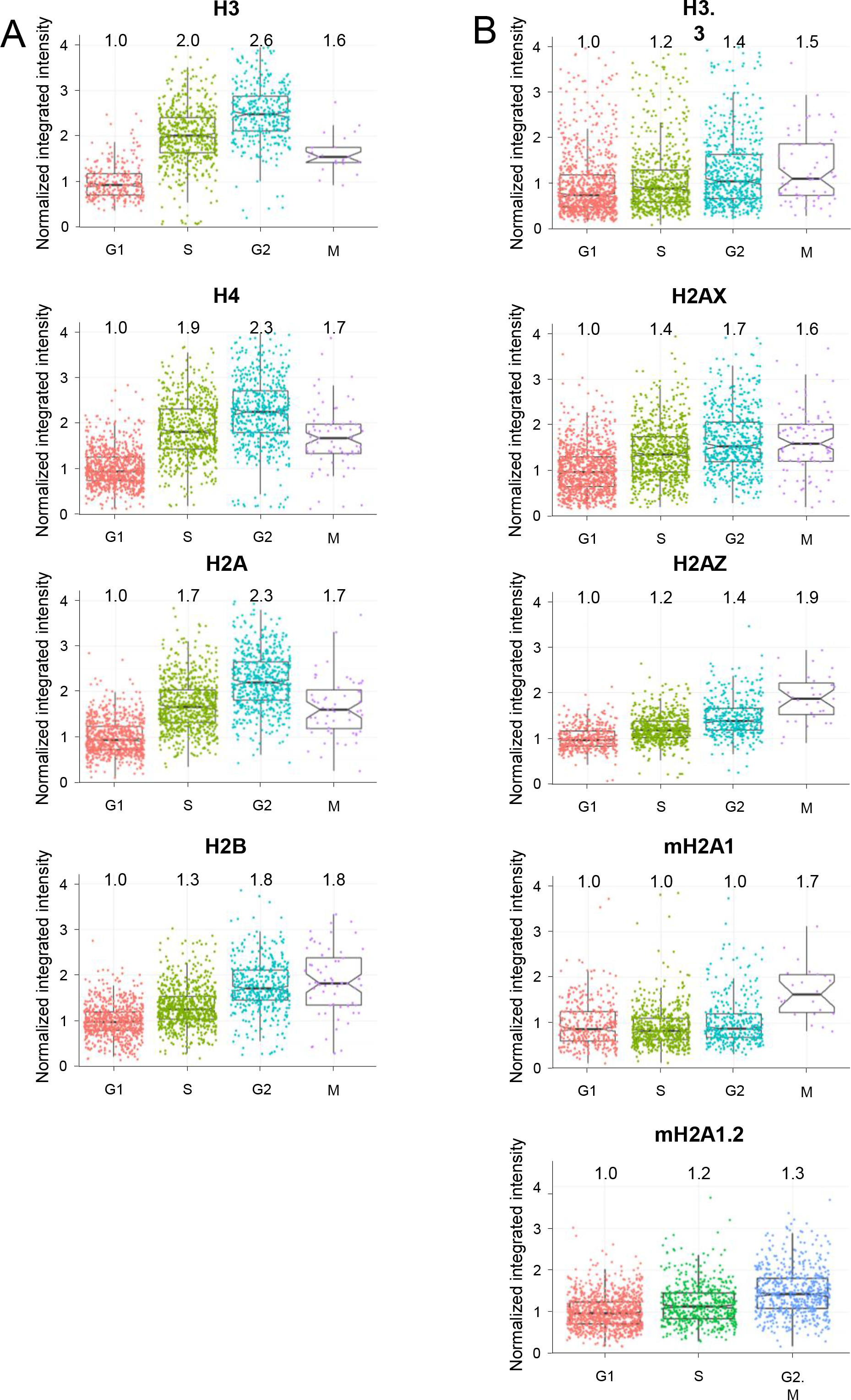
Single-cell levels of (A) core histones and (B) histone variants during the cell cycle. Each dot represents the level of the histone or histone modification of interest in a single cell. In box plots, the line corresponds to the median, notches represent the estimated 95% confidence interval (CI) for the median, the lower and upper hinges of the box plot indicate the 25th and 75th percentiles and the whiskers correspond to ± 1.5 * IQR of the hinge, where IQR is the interquartile range or distance between the first and third quartiles. The numbers above the box plots represent the means.

In contrast to the canonical histones, the synthesis of histone variants is uncoupled from DNA replication^47^. To characterize the cell cycle behavior of histone variants we applied HiHiMap to several histone variants (Fig. 3B). Levels of the H3 variant H3.3 and the H2A variant H2AZ levels only increased 1.3 ± 0.07- and 1.2 ± 0.13-fold from G1 to S phase and 1.3 ± 0.02- and 1.4 ± 0.02-fold to G2 before reaching a 1.7 ± 0.02- and 1.9 ± 0.09-fold increase in M phase (Fig. 3B). MacroH2A1 showed no increase from G1 to S and G2 phases (S/G1=1 ± 0.02;G2/G1=1 ± 0.03) followed by a 1.7 ± 0.12-fold increase between G2 and M. In contrast to other variants, H2AX, showed larger fluctuations from G1 to S and to G2 with 1.4 ± 0.02- and 1.7 ± 0.03-fold increases, respectively, followed by a slight decrease in mitosis (1.6 ± 0.07-fold decrease). Interestingly, the H2AX gene (*H2AFX*) contains features of both replication-dependent and replication-independent histone species, in that it is encoded by a small intronless gene and has the stem-loop structure that is characteristic of replication-linked histones^48^. We conclude that, in contrast to core histones, levels of several histone variants, except H2AX, do not follow cellular DNA levels.

### Characterization of cell-cycle fluctuations of histone modifications

Histones undergo extensive post-translational modifications^2,3^. The behavior of histone modifications during the cell cycle is relatively poorly characterized, in part due to the difficulty in generating cell-cycle staged cell populations for biochemical analysis and the heterogeneity of modifications amongst individual cells. We used HiHiMap to characterize the cell-cycle related fluctuations of 17 histone modifications and to determine the variability of modifications between individual cells in the same cell cycle phase.

Using a set of specific, previously characterized, antibodies (Table S1), we detect the rapid increase after DNA replication of several H3K4 methylation events. S/G1 ratios of mono-, di-and tri-methylation of H3K4 were increased 1.7 ± 0.03-, 1.6 ± 0.03- and 1.7 ± 0.02-fold, respectively, to reach a two-fold increase in G2 (H3K4-me1: ratio S/G1 = 2.1 ± 0.04; -me2: 1.9 ± 0.04; -me3: 2.1 ± 0.03) and then decreased in mitosis (ratios M/G1: 1.4 ± 0.01-fold, 1.6 ± 0.01-fold and 1.7 ± 0.09-fold for mono-, di-and tri-methylation, respectively (Fig. 4A). The level of H3K9me1, commonly associated with transcriptional activation^49^, followed the same pattern as H3K4 methylation during the cell cycle with its highest level in G2 phase (ratio S/G1=1.5 ± 0.02: ratio G2/G1= 2 ± 0.03; ratio M/G1=1.5 ± 0.09) (Fig. 4A). In contrast, the higher methylation states of H3K9, which are linked to transcriptional repression^50^, modestly increased from G1 to S (ratio S/G1=1.2 ± 0.01 for H3K9me2, ratio S/G1=1.3 ± 0.02 for H3K9me3) (Fig. 4A) and for H3K9me2 and H3K9me3 we observed 1.5 ± 0.02- and 1.4 ± 0.02-fold increases, respectively, in G2 phase and a 1.8 ± 0.07- and 2 ± 0.01--fold increase, respectively, in M phase compared to G1 (Fig. 4A). H3K27me3, one of the most prominent repressive histone marks^49,51^ increased by 1.6 ± 0.03- and 1.7 ± 0.04-fold in S and G2 phases, respectively, before dropping off to a 1.3 ± 0.08-fold increase in mitosis in comparison to G1 (Fig. 4A). H3K36me3, which is enriched in the transcribed regions of actives genes^52,53^, slightly increases during the cell cycle (ratios S/G1: 1.12 ± 0.02; G2/ G1:1.24 ± 0.02) before a 1.61 ± 0.01 increase in mitosis compared to G1 (Fig. 4A). Furthermore, mono-and di-methylated H4K20, involved in DNA replication and DNA repair^54^, modestly increased from G1 into S-phase (ratios S/G1:1.2 ± 0.03 for H4K20me1 and 1.1 ± 0.01 for H4K20me2) before increasing in G2 phase (ratios G2/G1:2.5 ± 0.08) for H4K20me1 and in mitosis for H4K20me2 (1.73 ± 0.09) (Fig. 4B).

**Figure 4.**
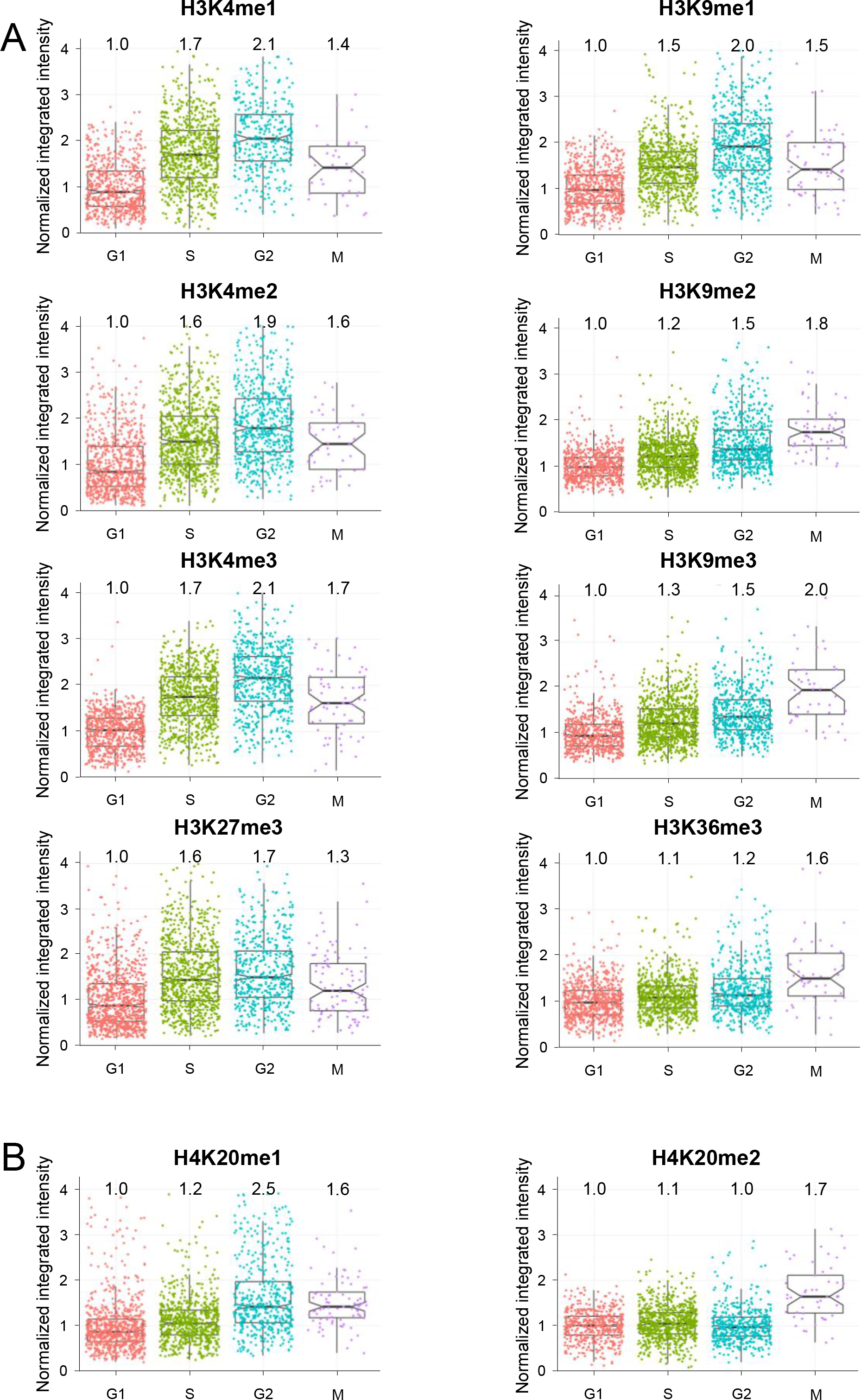
Single-cell levels of histone (A) H3 and (B) H4 methylation during the cell cycle. Each dot represents the level of the histone or histone modification of interest in a single cell. In box plots, the line corresponds to the median, notches represent the estimated 95% confidence interval (CI) for the median, the lower and upper hinges of the box plot indicate the 25th and 75th percentiles and the whiskers correspond to ± 1.5 * IQR of the hinge, where IQR is the interquartile range or distance between the first and third quartiles. The numbers above the box plots represent the means.

In contrast to the methylation levels, analysis by HiHiMap of H3K9Ac, H4K5Ac, and H3 and H4 global acetylation levels showed increases between G1 and S phases (ratio S/G1=1.52 ± 0.02 for H3K9Ac; 1.5 ± 0.02 for H4K5Ac; 1.5 ± 0.02 for H3Ac; 1.5 ± 0.02 for H4Ac), and even further after DNA synthesis leading to a two-fold ratio between G2 and G1 phases (ratio G2/G1=2 ± 0.025 for H3K9Ac;2 ± 0.037 for H4K5Ac;2.1 ± 0.04 for H3Ac;1.9 ± 0.03 for H4Ac) followed by a slight decrease in mitosis (ratio M/G1=1.86 ± 0.07 for H3K9Ac; 1.73 ± 0.01 for H4K5Ac; 1.7 ± 0.01 for H3Ac; 1.6 ± 0.1 for H4Ac) (Fig. 5A, B). These observations delineate the cell-cycle behavior of a comprehensive sets of histone modifications at the single-cell level.

**Figure 5.**
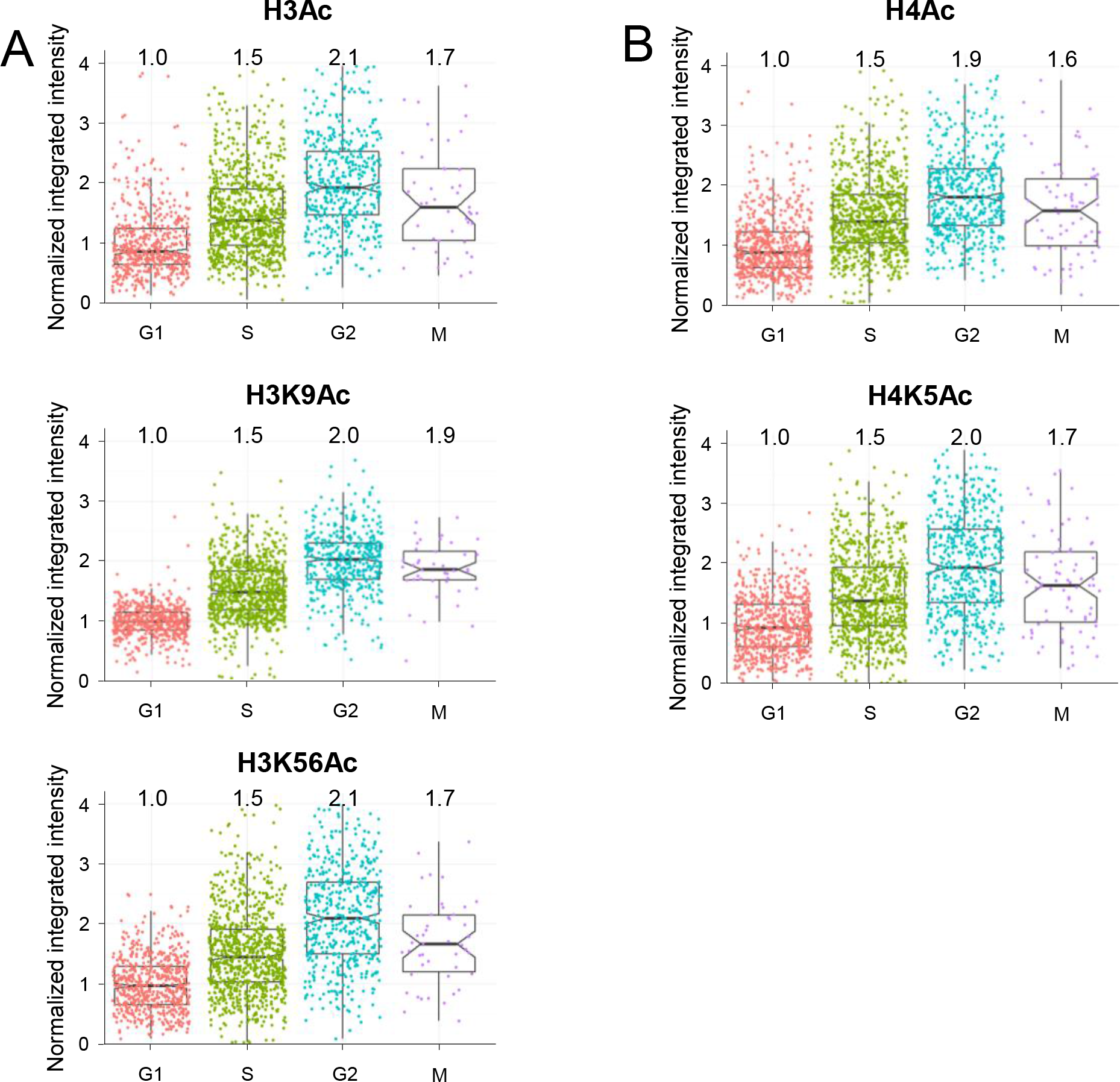
Single-cell levels of histone (A) H3 and (B) H4 acetylation levels during the cell cycle. Each dot represents the level of the histone or histone modification of interest in a single cell. In box plots, the lined corresponds to the median, notches represent the estimated 95% confidence interval (CI) for the median, the lower and upper hinges of the box plot indicate the 25th and 75th percentiles and the whiskers correspond to ± 1.5 * IQR of the hinge, where IQR is the interquartile range or distance between the first and third quartiles. The numbers above the box plots represent the means.

### Identification of dysregulated histone and histone post-translational modifications during oncogenic transformation

To apply HiHiMap to a biological problem, we took advantage of the capacity of HiHiMap for multiplexing to systematically analyze histones and histone modifications during oncogenic transformation. To generate oncogenically transformed cells we *in vitro*-transformed primary human skin fibroblasts from three individuals by introduction of hTERT, H-RasV12 and SV40 Large-T (LT) and Small-T (ST) antigens as previously described^33–36^ (Fig. S3). We confirmed the presence of the three transgenes at the RNA (Fig. S3A) and protein (Fig. S3B) levels and their ability to form colonies in soft agar at high rates (Fig. S3C). Comparative profiling of the cell cycle distribution of primary, immortalized and transformed cells by DAPI staining revealed the expected high levels of aneuploidy in the transformed cells with the shift of the cell cycle profile towards higher levels of DAPI integrated intensity (Fig. S4, 5). These results were confirmed by flow cytometry (Fig. S4).

To account for the tetraploid nature of transformed cells, the DAPI thresholds to stage cells were modified and diploid G2 cells were defined as cells with DAPI intensity of 1.75 – 2.25 and cyclin A positive, and tetraploid cells in G1 phase as DAPI intensity of 1.75 – 2.25 and cyclin A negative. Tetraploid cells in S phase were defined as DAPI intensity of 2.25 – 2.75 and cyclin A positive and diploid cells in G2, as a DAPI intensity of 1.75 – 2.25 and cyclin A positive (Fig. S6). The ploidy status of the transformed cells was confirmed by flow cytometry to correctly assign cell cycle stages in these samples (Fig. S5). Correspondingly, transformed cells exhibited higher background levels of cyclin A and thresholds were adjusted accordingly (Fig. S7). To account for the variability in genetic material between individual cells, histone levels were normalized to the amount of DNA in each cell.

The values of 22 histone variants and modifications of primary, immortalized and transformed cells in each cell cycle stage are represented as a heat-map in Figure 6. We normalized the levels of core histones and histone variants to the DNA content (Fig. 6C). We observed an increase of 2.1 ± 0.02-, 1.5 ± 0.07-, 1.4 ± 0.04- and 2.7 ± 0.03-fold in the level of H2AX variant in G1, S, G2 and M, respectively, between the primary and transformed cells (p < 10^-14^ for each cell cycle stage, Student's t-test) and an increase of 2.6 ± 0.03-, 1.7 ± 0.05, 1.8 ± 0.03- and 3.3 ± 0.08-fold in the level of this variant between the immortalized cells and their transformed counterparts in G1, S, G2 and M (p < 10^-16^, Student’s t-test), respectively (Fig. 6C, Fig. 7A). In contrast, we observed a slight decrease of 0.81 ± 0.04-(p < 10^-16^), 0.87 ± 0.15-(p = 0.07), 0.81 ± 0.09-(p = 4.59 × 10^-5^) and 0.82 ± 0.11-fold (p = 0.001, Student's t-test) in the levels of H2AX between the primary and immortalized cells in G1, S, G2 and M phases, respectively. Representative results for a single cell line (AG06310) are shown and all results were confirmed in three independent experiments in HDFs from three individuals (Fig. 6, Fig. S8, Fig. S9).

**Figure 6.**
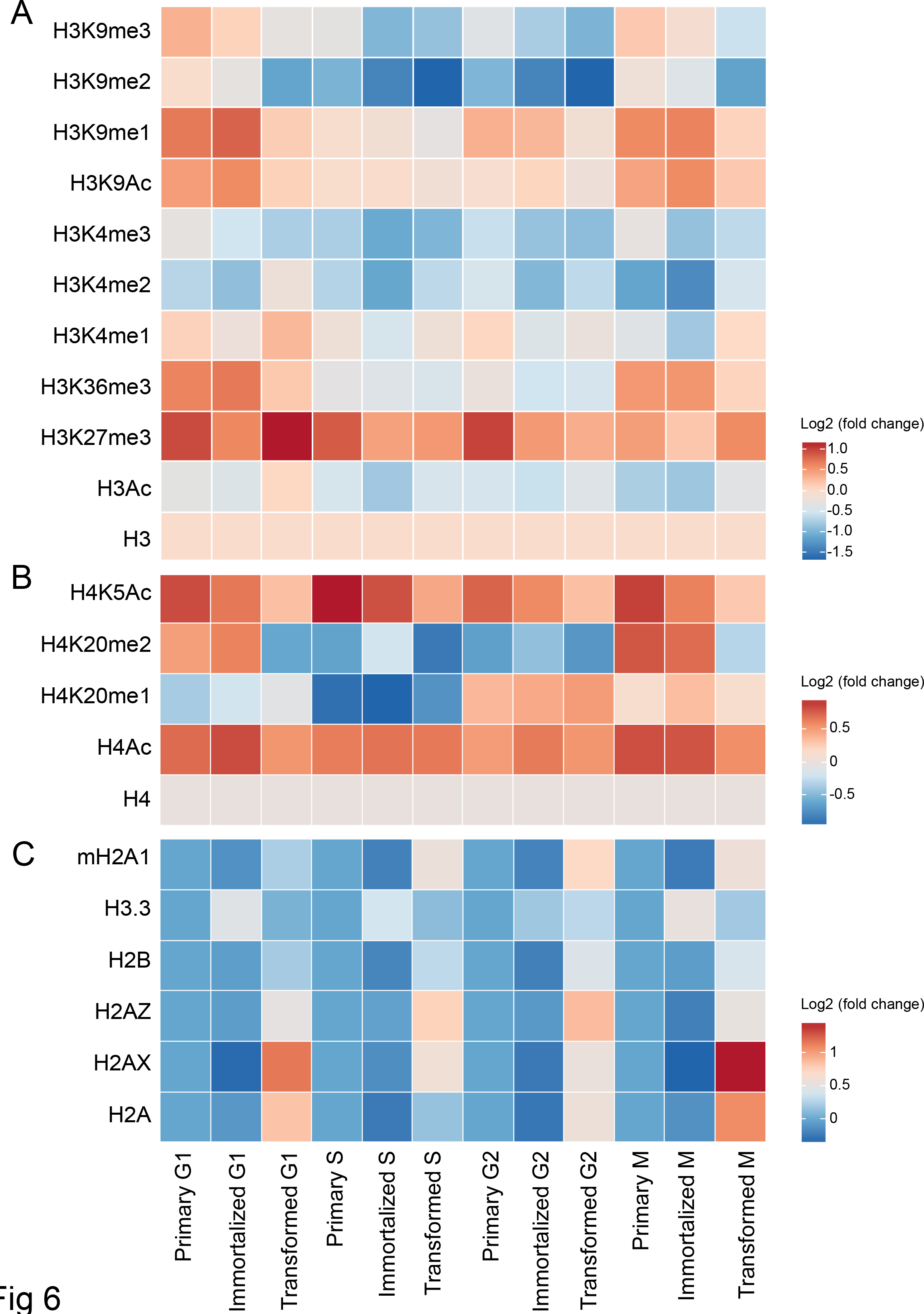
Heat-maps of changes in histone and histone PTM levels during oncogenic transformation of human skin fibroblasts in the different cell cycle phases. Representation of the fold changes in (A) H3 modification levels normalized to H3 levels, (B) H4 modification levels normalized to H4 levels and (C) histone and histone variant levels in the primary, immortalized and transformed cells in AG06310 cells in G1, S, G2 and M phases after normalization to the level of their primary counterparts in the same cell cycle phase.

**Figure 7.**
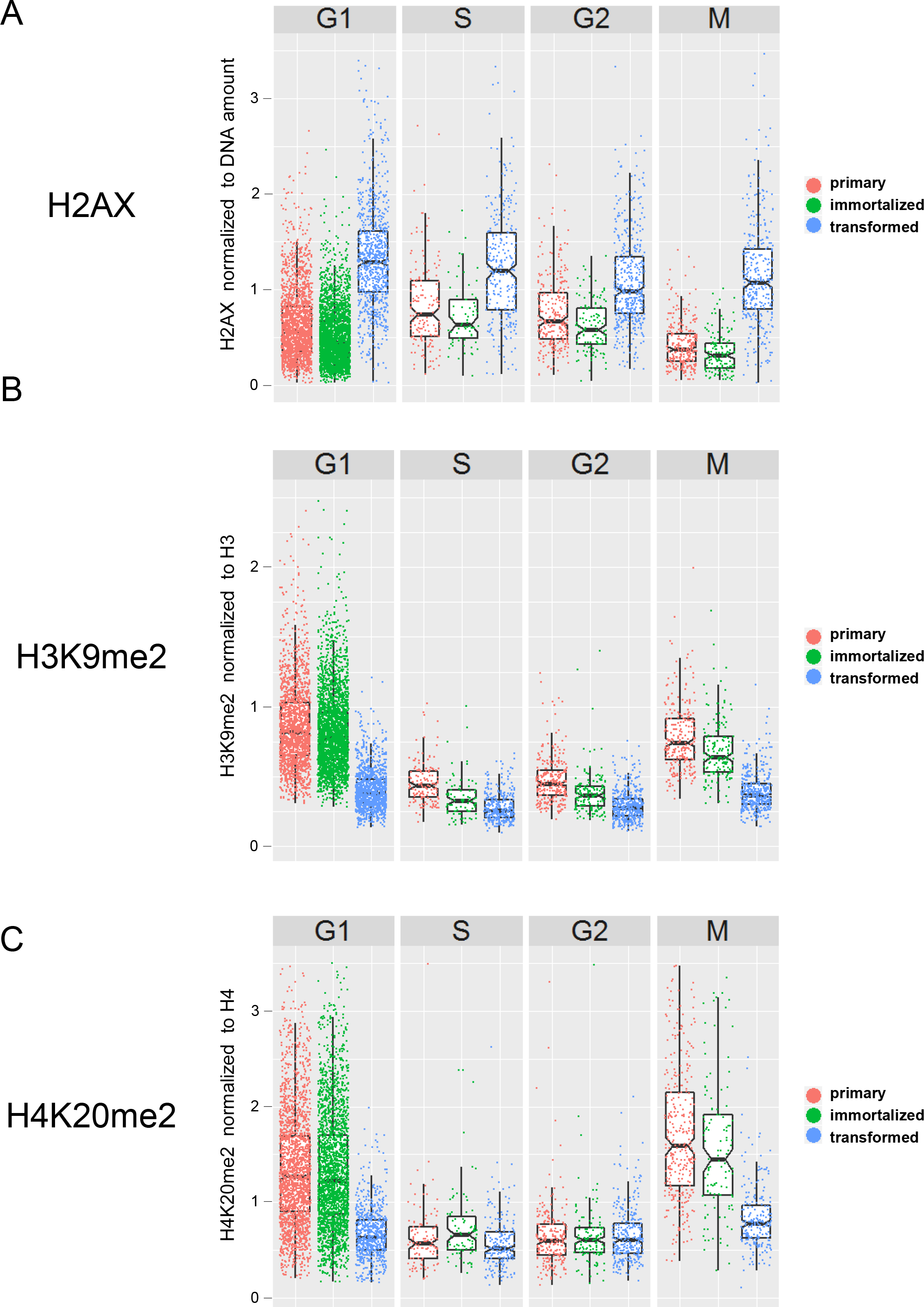
Single-cell intensity levels of (A) histone H2AX normalized to DNA amount, (B) H3K9me2 normalized to H3 levels and (C) H4K20me2 normalized to H4 levels in primary, immortalized and transformed cells in AG06310 cells in G1, S, G2 and M phases. Each dot represents the level of the histone or histone modification of interest in a single cell. In box plots, the lined corresponds to the median, notches represent the estimated 95% confidence interval (CI) for the median, the lower and upper hinges of the box plot indicate the 25th and 75th percentiles and the whiskers correspond to ± 1.5 * IQR of the hinge, where IQR is the interquartile range or distance between the first and third quartiles.

For analysis of modifications of histones H3 and H4, we normalized their levels to the levels of the corresponding core histone to account for the observed aneuploidy in individual cells and to account for the variations in the levels of core histones between samples (Fig. 6A and B). After normalization to H3, H3K9me2 was decreased by 0.45 ± 0.03-(p < 10^-16^), 0.44 ± 0.47-(p=0.055), 0.61 ± 0.05-(p < 10^-16^,), 0.49 ± 0.06-fold (p < 10^-16^) between primary and transformed cells and by 0.47 ± 0.03-(p < 10^-16^,), 0.66 ± 0.047-(p=0.5×10^-4^), 0.73 ± 0.05-(p=1.9×10^-14^) and 0.5 ± 0.06-fold (p < 10^-16^) between immortalized and transformed cells in G1, S, G2 and M phases, respectively (Fig. 6A, Fig. 7B). The same observation was made in HDFs from another individual (AG04551, see Materials and Methods; Fig. S9).

We observed a statistically significant decrease of 0.6 ± 0.04-(p < 10^-16^) and 0.53 ± 0.12-fold (p < 10^-16^) in the level of H4K20me2 normalized to H4 between primary and transformed cells as well as a decrease of 0.6 ± 0.04-(p < 10^-16^) and 0.59 ± 0.12-fold (p=1.33×10^-5^) between immortalized and transformed cells in G1 and M phases, respectively (Fig. 6B and Fig. 7C). All results were confirmed in three independent experiments and similar results were observed in HDFs from three individuals (Fig. 6, Fig. S8, Fig. S9).

HiHiMap was also able to detect differences in the behavior of individual histones and histone modifications across the cell cycle (Fig. 8). For this analysis values were normalized to the G1 values for each modification. The values of 22 histone variants and modifications across the cell cycle of primary, immortalized and transformed cells are represented as a heat-map in Figure 8. For example, the level of H4K20me2 normalized to H4 was relatively stable during the cell cycle in transformed cells with a 0.84 ± 0.05-, 0.9 ± 0.04- and 1.1 ± 0.04-fold change in S, G2 and M phases compared to G1 (p-values: 3.4×10^-4^, 1.3×10^-3^ and 1.1×10^-2^). However, a decrease of 0.47 ± 0.04- and 0.48 ± 0.04 fold in S and G2 phases followed by an increase of 1.3 ± 0.01-fold in mitosis compared to G1 were observed in primary cells (p < 10^-16^ for all S vs. G1, G2 vs. G1 and M vs. G1 comparisons). A decrease of 0.57 ± 0.03- and 0.54 ± 0.04-fold for H4K20me2 in S and G2 phases followed by a 1.15 ± 0.02-fold increase in M were also observed in immortalized cells (p-values: 4.4×10^-15^, 6.5×10^-16^ and 5.4×10^-3^) (Fig. 7C, Fig. 8B). Similarly, the level of H2AX is stable during the cell cycle in transformed cells with a 1.3 ± 0.03-, 1.6 ± 0.02- and 1.7 ± 0.06-fold change in S, G2 and M phases compared to G1 (p-values: 7.7×10^-14^ in S and p<10^-16^ in G2 and M), but fluctuates by 1.9 ± 0.02-(p<10^-16^) and 2 ± 0.01-fold (2.5x10^-12^) in S, by 2.4 ± 0.01-(p<10^-16^) and 2.4 ± 0.014-fold in G2 (p<10^-16^), and by 1.3 ± 0.02-(p=2.1×10^-13^) and 1.4 ± 0.02-fold (p=1.96×10^-6^) in M phase compared to G1, respectively, in primary and immortalized cells (Fig. 7C, Fig. 8C). In contrast, the level of H2AZ varies considerably in S and G2 phases of transformed cells (1.6 ± 0.02-fold, p<10^-16^ in S and 2 ± 0.01-fold, p<10-16 in G2), and varies less in their primary (1.4 ± 0.01-fold, p=7.1×10^-11^ in S; 1.6 ± 0.01-fold, p<10^-16^ in G2) and immortalized (1.3 ± 0.01-fold, p=7.6×10^-6^ in S; 1.5 ± 0.01-fold, p<10^-16^ in G2) counterparts (Fig. 7C, Fig. 8C). All results were confirmed in three independent experiments and similar results were observed in HDFs from three individuals (Fig. 7, Fig. 8, Fig. S10, Fig. S11). These results point to considerable effects on histone and histone modifications during oncogenic transformation.

**Figure 8.**
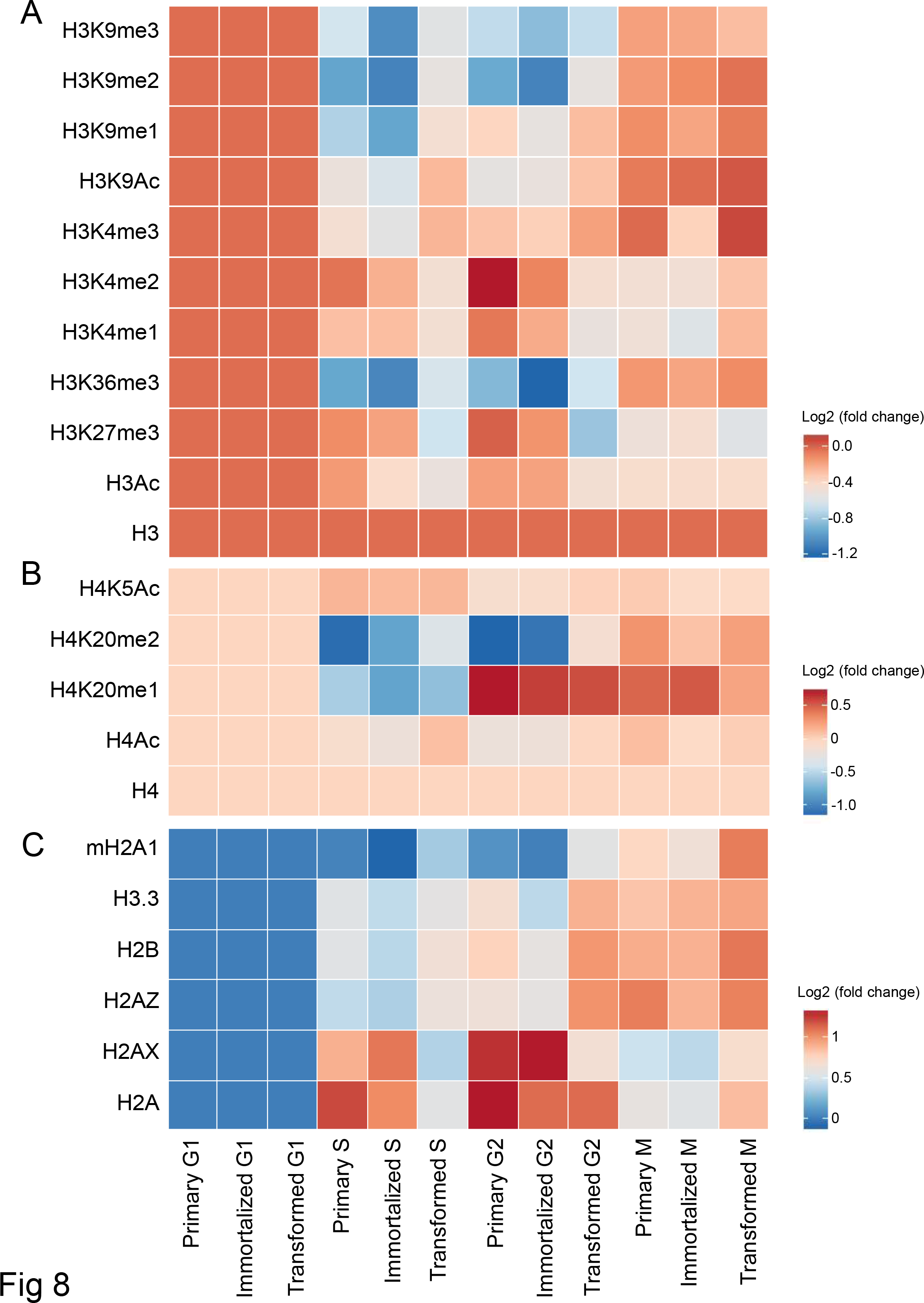
Heat-maps of changes in histone and histone PTM levels during the cell cycle detected by high-throughput confocal microscopy. Representation of the fold changes in (A) H3 modification levels normalized to H3 levels, (B) H4 modification levels normalized to H4 levels and (C) histone and histone variant levels of primary, immortalized and transformed cells in AG06310 cells in G1, S, G2 and M phases after normalization to their respective level in G1.

## DISCUSSION

We have developed an imaging-based method, termed HiHiMap, to quantitatively measure histone and histone PTM levels throughout the cell cycle at the single-cell level using high-throughput microscopy. For validation we show that HiHiMap is able to detect several expected changes in histones and histone modifications through the cell cycle, we demonstrate proof-of-principle by measurement of a comprehensive set of 26 histones and histone modifications and we demonstrate its ability for discovery by identification of changes in a wide range of histone modifications during oncogenic transformation.

HiHiMap overcomes several limitations of traditional methods, such as Western blotting and SILAC combined with mass spectrometry, to determine histone and histone modification levels across the cell cycle. HiHiMap does not require any cell synchronization, which often results in impure populations, nor does it necessitate *in vivo* protein labeling steps, which may interfere with protein function. HiHiMap also provides practical advantages in that it is a rapid and easy method to study a wide range of histones and histone modifications in a fully automated fashion, enabling comprehensive analysis of a large number of histones and histone modifications in a single experiment. In addition, considerably lower numbers of cells are needed for HiHiMap (3,000-5,000 cells per sample) than for the other methods (typically 10^5^-10^6^ cells per sample), which is particularly important when working with patient samples. Furthermore, HiHiMap allows visualization of the distribution of histones and PTM in individual nuclei, which enables correlations of histone and histone modification levels relative to cellular features, for example the formation of senescence associated heterochromatin foci in individual cells in the population^55,56^ or, as shown here, relative to the ploidy level of individual cells, a feature important in the analysis of cancer samples^57^. Most importantly, HiHiMap is a single-cell analysis method and, in contrast to biochemical methods which use bulk analysis and generate population averages, it measures histone and histone PTM in single cells whose position in the cell cycle has been determined. The method thus allows the study of the behavior of subpopulations of cells and of the variability within a population. For example, we find much greater variability in the single-cell distribution of histone and PTM levels in transformed cells than within the primary or immortalized populations (Fig. 7A). In addition, pooling of single cell data across the typically 1,000-2,000 analyzed cells also allows derivation of population averages, making the data comparable to biochemical datasets.

A limitation of HiHiMap is that it cannot distinguish between free and bound histones in the nucleus as well as between the same modifications on multiple isoforms of core histones. In addition, HiHiMap also does not distinguish pre-existing from newly deposited histones or histone modifications after DNA replication. Furthermore, the method requires use of antibodies for detection. However, the abundance of well-characterized antibodies against a wide array of histones and histone modifications in various species lessens this concern.

We validated HiHiMap using several replication-dependent histones and demonstrate accurate detection of their expected increases. As proof-of-principle for its application as a discovery tool, we investigated global changes of histone and histone PTM levels during oncogenic transformation. In an established cell-based transformation model we find a number of changes in histones and histone modifications over the course of oncogenic transformation, pointing to potential biological mechanisms. Of significance, we find reduction of H3K9me2, which is known to characterize condensed heterochromatin, and thus gene inactivation^49,58,59^, and has recently been shown to be decreased in an *in vitro* tumor transformation model of human primary mammary cell and in breast cancer tissues^60^. Similarly, global levels of H3K9 dimethylation are significantly lower in bladder cancer than in normal urothelial tissue, and levels in muscle invasive bladder cancer are lower compared to non-muscle invasive bladder cancer^61^. In prostate^62^ and kidney cancers^16^ as well as in clear cell renal cell carcinoma^63^, lower levels of H3K9me2 have been associated with disease outcome and predict poor prognosis. We also found a decrease in H4K20 dimethylation, the most abundant methylation mark on histone H4 in proliferating cells^29^, which has been linked to DNA damage and DNA repair^64^ and genomic instability^54^. In agreement with our results, hypomethylation of H4K20me2 has been shown in metastatic and castration-resistant prostate cancer compared to normal prostate tissue^65^ and in several other cancers including bladder^66^and liver^67^ tumors. Finally, we found increased levels of H2AX in transformed cells compared to their primary and immortalized counterparts. The histone variant, H2AX, has been implicated in tumor development by virtue of frequent mutations and deletions of chromosome region 11q23, which contains the H2AX gene, in a large number of human cancers, especially in hematopoietic malignancies^70,71^ and head and neck squamous cell carcinoma^72^ and H2AX knockout mice show increased genomic instability and a higher risk of developing cancers^73^. In addition, phosphorylation of H2AX plays a key role in the DNA damage response^74^ and its levels are increased in many cancer cells^75–78^.

In summary, we have developed a versatile method to measure in individual cells the levels of multiple histone and histone PTMs throughout the cell cycle. We validated the method and applied it to identify changes in histones and histone modifications during oncogenic transformation. We anticipate that HiHiMap will be a useful tool to study the regulation of histone and histone PTM levels through the cell cycle in a wide range of physiological and pathological conditions.

## ACKNOWLEDGEMENTS

We thank Sigal Shachar for technical help with the Opera and members of the Misteli laboratory for helpful discussions and comments. This research was supported by the Intramural Research Program of the National Institutes of Health (NIH), National Cancer Institute, and Center for Cancer Research.

## Supplementary figure legends

**Table S1.** List of antibodies used.

**Figure S1.** Cell cycle staging using DAPI and cyclin A staining. Representative confocal 40X images of cyclin A and DAPI staining acquired in two channels in immortalized human dermal fibroblasts CRL1474-hTERT. Scale bar, 10 μm. The histograms of DAPI and cyclin A intensities in the cell population are calculated. Using the DNA content (DAPI intensity), cell cycle profiles are generated and visually selected cutoffs identify cells in the different cell cycle phases G1, S, G2/M. The cutoffs are determined in the cell cycle profiles after normalization of the first peak at 1 to be able to apply similar cutoffs for different wells of a same plate. Cyclin A positive cells are also identified by applying a visually selected cutoff. By using the double staining cyclin A/DAPI and their cutoffs, every individual cell is associated to a cell cycle stage.

**Figure S2.** Reproducibility of HiHiMap. Differences in the level of core histone H4 through the cell cycle are technically and biologically reproducible. The experiment was performed three independent times and every experiment consisted of two technical replicates.

**Figure S3.** *In vitro*-transformation of human primary skin fibroblasts from 3 healthy individuals (AG06310, AG06289 and AG04551). (A) Expression of SV40 LT and H-Ras proteins in transformed fibroblasts were detected by Western blot. The anti-Ras antibody recognizes K-, N-, and both wild-type and mutated H-Ras. GAPDH was used as loading control. (B) Quantitative RT-PCR measuring expression levels of *HTERT, SV40 LT* and *HRASV12* in primary, hTERT-immortalized and transformed hRS (*h*TERT+*R*asV12+*S*V40) cells. Primers for *HRASV12* do not detect endogenous *HRAS.* Values are normalized to the housekeeping gene *GAPDH* and represent mean ± s. d. from two replicates. (C) Soft-agar assay using hTERT-immortalized and hRS-transformed cells. Scale bar, 8.89 μm.

**Figure S4.** Comparison of cell cycle profiles obtained by traditional, population-based DNA content analysis by flow cytometry (propidium iodide staining, left) and by single-cell confocal high-throughput microscopy (DAPI staining, right). Cell cycle analysis of primary, immortalized and transformed cells from three samples: (A) AG06310, (B) AG06289 and (C) AG06310 by imaging and flow cytometry.

**Figure S5.** Aneuploidy status of transformed samples from three different individuals determined by flow cytometry. Cell cycle profiles of primary, immortalized, transformed and a mix of primary and transformed cells from three different samples: (A) AG06310, (B) AG06289 and (C) AG06310 obtained by population-based DNA content analysis by flow cytometry (propidium iodide staining).

**Figure S6.** Adaptation of the high-throughput histone mapping (HiHiMap) method for the study of oncogenically transformed samples. The ploidy status of cells is affected by oncogenic transformation (see Figure S5). To simplify the representation of the results, since the main population corresponds to the tetraploid population, only their levels of histones and histone PTM through the cell cycle is represented. The levels of core histones and histone variants are normalized to the DNA amount and the levels of histone modifications normalized to the corresponding core histone.

**Figure S7.** Determination of cyclin A thresholds in primary, immortalized and transformed samples from the same individual. Cyclin A thresholds are visually determined using the background intensity level corresponding to the intensity level without cyclin A staining (left). The populations of cyclin A positive and negative cells are identified based on these thresholds that are different between samples.

**Figure S8.** Heat-maps summarizing the changes in histone and histone PTM levels during oncogenic transformation of human skin fibroblasts in the different cell cycle phases detected by high-throughput confocal microscopy. Representation of the fold changes in (A) H3 modification levels normalized to H3, (B) H4 modification levels normalized to H4 and (C) histone and histone variant levels in the primary, immortalized and transformed cells in AG06289 cells in G1, S, G2 and M phases after normalization to the level of their primary counterparts in the same cell cycle phase.

**Figure S9.** Heat-maps of the changes in histone and histone PTM levels during oncogenic transformation of human skin fibroblasts in the different cell cycle phases detected by high-throughput confocal microscopy. Representation of the fold changes in (A) H3 modification levels normalized to H3, (B) H4 modification levels normalized to H4 and (C) histone and histone variant levels in the primary, immortalized and transformed cells in AG04551 cells in G1, S, G2 and M phases after normalization to the level of their primary counterparts in the same cell cycle phase.

**Figure S10.** Heat-maps of changes in histone and histone PTM levels during the cell cycle detected by high-throughput confocal microscopy. Representation of the fold changes in (A) H3 modification levels normalized to H3, (B) H4 modification levels normalized to H4 and (C) histone and histone variant levels of primary, immortalized and transformed cells in AG06289 cells in G1, S, G2 and M phases after normalization to their respective level in G1.

**Figure S11.** Heat-maps of changes in histone and histone PTM levels during the cell cycle detected by high-throughput confocal microscopy. Representation of the fold changes in (A) H3 modification levels normalized to H3, (B) H4 modification levels normalized to H4 and (C) histone and histone variant levels of primary, immortalized and transformed cells in AG04551 cells in G1, S, G2 and M phases after normalization to their respective level in G1.

